# Indirect genetic effects clarify how traits can evolve even when fitness does not

**DOI:** 10.1101/458695

**Authors:** David N. Fisher, Andrew G. McAdam

## Abstract

There are many situations in nature where we expect traits to evolve but not necessarily for mean fitness to increase. However, these scenarios are hard to reconcile simultaneously with Fisher’s Fundamental Theorem of Natural Selection and the Price identity. The consideration of indirect genetic effects on fitness reconciles these fundamental theorems with the observation that traits sometimes evolve without any adaptation, by explicitly considering the correlated evolution of the social environment, which is a form of transmission bias. While transmission bias in the Price identity is often assumed to be absent, here we show that explicitly considering indirect genetic effects as a form of transmission bias for fitness has several benefits: 1) it makes clear how traits can evolve while mean fitness remains stationary, 2) it reconciles the fundamental theorem of natural selection with the evolution of maladaptation, 3) it explicitly includes density-dependent fitness through negative social effects that depend on the number of interacting conspecifics, and 4) its allows mean fitness to evolve even when direct genetic variance in fitness is zero, if related individuals interact and/or if there is multilevel selection. In summary, considering fitness in the context of indirect genetic effects aligns important theorems of natural selection with many situations observed in nature and provides a useful lens through which we might better understand evolution and adaptation.

## Fundamental theorems of evolution and adaptation

R. A. Fisher’s “fundamental theorem of natural selection” (FTNS) is one of the most famous, and still widely debated ideas in evolutionary biology (Fisher 1930). Following careful reevaluation by G. R. Price, it is generally understood that Fisher’s FTNS should be understood as: In any population at any time, the rate of change of fitness ascribable to natural selection is equal to its additive genetic variance at that time (Price 1972). This is:

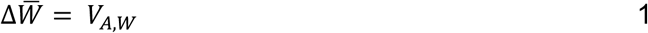

Where Δ*W̅* refers to the change in mean fitness from one generation to the next caused by natural selection, and *V_A,W_* is the additive genetic variance in fitness. Fitness here is “lifetime breeding success” or similar, i.e. an absolute value, as Fisher related it to population growth (Fisher 1930). Recent commentators have concluded that the FTNS is essentially true, and in the way Fisher meant it (Bijma 2010a; Grafen 2015; Birch 2016). Therefore, when *V_A,W_* > 0, natural selection is causing mean fitness to increase. Note that mean fitness may also be increased or decreased by changes in the environment, hence the change ascribable to natural selection may not be equal to observed changes in fitness, but for our purposes here we assume a constant abiotic environment.

Independently derived, but fundamentally linked (Queller 2017), is the Price identity (Price 1970; hereafter the PI, note a similar expression, but lacking the second term, was derived earlier by A. A. Robertson 1966):

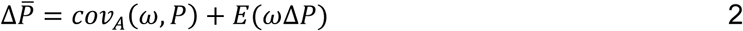

Where Δ*P̅* refers to the change in mean phenotypic trait value from one generation to the next, and *cov_A_*(*ω,P*) to the additive genetic covariance between individuals’ relative fitness (*ω*, equal to 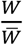) and some phenotype (*P*) and *E*(*ω*Δ*P*)) is the change in mean phenotype between parents and offspring, which could be caused by a bias in meiosis or fertilisation, or by changes in the environment, which is referred to as “transmission bias”. This simple but powerful expression for the expected change in phenotypes states that for evolution to occur, there must be a genetic covariance between relative fitness and the trait in question.

In typical treatments of trait evolution based on the Price identity, researchers assume that the transmission bias is equal to zero, which gives Robertson’s expression for the evolution of traits (Robertson 1966). We do not contend this is incorrect, but we highlight later that a portion of the change partitioned to transmission bias will in fact often have an additive genetic basis, and therefore considering it explicitly is essential to understand evolutionary trajectories in some cases. Otherwise, we assume a constant abiotic environment throughout. Although it is not always appreciated, the PI implies that *for any trait to evolve there must be non-zero additive genetic variance in fitness*, otherwise the genetic covariance is undefined and evolution does not proceed (Morrissey et al. 2010; Shaw and Shaw 2014).

The PI therefore makes clear that if any trait is evolving, there must be genetic variance in fitness. Further, if there is genetic variance in fitness (*V_AW_* > 0), then according to the FTNS mean fitness must be increasing (Δ*W̅* > 0). Conversely, if mean fitness is not being increased by natural selection (Δ*W̅* = 0) then genetic variance in fitness must be zero (*V_A,W_* = 0) and so no trait can evolve. The combination of Fisher’s FTNS and the PI, therefore, lead to the following statements:

> *“If a trait is evolving by natural selection, there must be genetic variance in fitness, and so mean fitness is evolving”*
>
> and
>
> *“If a population’s mean fitness is not evolving, then additive genetic variance in fitness must be zero, so no trait can evolve as a result of natural selection”*

We refer to situations where some trait is evolving in response to natural selection as “evolution by natural selection”, while we refer to situations where mean fitness is increasing by evolution as “adaptation”. Taking the FTNS and the PI together implies evolution by natural selection is *always* associated with adaptation. There are, of course, may ways in which changes in the environment might cause mean fitness to remain stationary or decline, but here we consider scenarios where the external environment remains constant.

In contradiction with these statements derived from the FTNS and PI, we clearly observe situations in nature where evolution occurs, but adaptation does not (Fisher 1941; Cooke et al. 1990; Frank and Slatkin 1992; Wolf et al. 2008). An example of this is that males with larger weapons, or preferred sexual displays, are expected to sire more offspring than their less well-endowed conspecifics. If these sexually selected male traits are heritable, we would expect the mean trait to change across generations; we therefore have a genetic covariance between the trait and fitness that is greater than zero. If so, there must be additive genetic variance in fitness, and so Fisher’s FTNS predicts that mean fitness ought to evolve (Δ*W̅* > 0). However, in reality there is no expectation that the *total* amount of reproductive success in the population will evolve, i.e. in this situation we would not expect females to start having more offspring, and so mean fitness is not expected to change. Therefore, no adaption is occurring, and following Fisher’s FTNS, genetic variance in fitness ought to be zero (*V_A,W_* = 0). Following the PI, evolution should then be impossible, yet we clearly expect the weapons or the display trait to evolve if they are heritable. This scenario also applies to any example of “soft” selection, where selection occurs among-individuals, but does not lead to the mean reproductive output increasing (as opposed to “hard” selection, where selection does lead to an increase in mean fitness Wallace 1975). So how can we explain the action of sexual and soft selection, given that the FTNS and the PI are true? To put it another way, when mean fitness is not evolving, do we really expect all evolution to cease?

Furthermore, we can observe situations where trait evolution (requiring non-zero *V_A,W_*) leads to reduced rather than increased fitness (“maladaptation”, distinct from situations where mean fitness is reduced purely by a change in the environment; Crespi 2000; Rogalski 2017). For example, *Agelenopsis aperta* spiders in riparian zones show suboptimal foraging and anti-predator behaviours compared to grassland populations, despite the riparian habitat being available for at least 100 years (Riechert 1993). The FTNS suggests that, as *V_A,W_* cannot be less than zero, Δ*W̅* cannot be negative. Therefore, the FTNS seems incompatible with observations of the evolution of maladaptation.

## Social interactions as part of the environment

This paradox can be resolved by revisiting an element of the PI that is typically set aside: the transmission bias. A transmission bias occurs when the mean phenotype of offspring and parents differ, but not due to evolutionary change (Frank 2012). Typical examples are when meiosis or fertilisation are not random with respect to the genes of interest, or when the environment has changed in some way, and organisms’ traits depend on this environment. Fisher too had a term for when phenotypes differ across generations due to environmental change (“environmental deterioration”), and noted that it would typically act to reduce mean fitness, which otherwise would continually increase (Fisher 1930). Fisher and others considered the competitiveness of conspecifics to be a key part of the environment (Fisher 1930; Cooke et al. 1990; Frank and Slatkin 1992). Importantly, this “social environment” is partly genetic in basis (as social traits will be partly heritable like any other trait) and so can evolve (Griffing 1967; Moore et al. 1997). Hence a possible source of transmission bias and environmental deterioration with limitless potential to continually change is the social environment. Here we contend that not only can the social environment evolve, but that with respect to many situations there are strong reasons to believe that *the social environment must evolve*. Explicitly considering the evolution of the social environment and its influence on the evolution of transmission bias allows trait evolution and adaptation to become dissociated.

As an example of how the evolution of the social environment will dissociate trait evolution from adaptation, we can consider the evolution of the ability to win contests for dominance in a dyadic interaction, such as when two stags square off to determine who is the strongest. Winning contests generally gives fitness benefits, and the propensity to win contests is also often heritable (Wilson et al. 2009, 2011), so we would expect the mean tendency to win such interactions to evolve. However, following Wilson and colleagues (2009, 2011; 2014), a “common-sense” approach sees this is impossible, because in every dominance interaction, there must be one winner and one loser, and hence the mean outcome in a dyadic contest is constrained to remain half winning and half losing in each generation. This is analogous to a situation where mean reproductive output cannot evolve, for instance when it is constrained at the population level by resource availability (be that food, territory space, or total offspring production of females in the case of sexual selection) even though increased reproductive output is always expected to be favoured by fecundity selection (Cooke et al. 1990; Frank and Slatkin 1992).

Common sense and models for micro-evolutionary change are reconciled by appreciating that individuals possess genetic effects for their *opponent’s* ability to win the dominance interaction (Wilson et al. 2009, 2011; Wilson 2014). In a zero-sum contest, where one individual’s success directly detracts from their competitor’s success, genes that enhance an individual’s chance of winning a contest necessarily reduce their opponent’s chance of winning. As these genes will be selected for, the propensity to win evolves, but so too does the propensity for others to lose as a correlated response. As opponents are drawn from the same population, contests for dominance in the next generation are now with more competitive opponents, i.e. the environment has evolved to become more competitive at the same time (Wilson 2014). This leads to no change in mean phenotype overall. This has been termed the evolution of environmental deterioration as the environment the trait (winning contests) is being expressed in has deteriorated (i.e. it has become more difficult to express the trait; Fisher 1930). Crucially, there is still direct genetic variance in the population for dominance, and so breeding values for it will increase over time. As such, traits correlated with direct breeding values for the ability to win contests, such as weapon size, will still evolve.

We can consider the importance of the evolution of the social environment to trait evolution and adaptation in general by considering a quantitative genetic model of trait evolution that considers indirect genetic effects (IGEs). Indirect genetic effects occur when the phenotype of one individual is affected by the genotype of another individual (Moore et al. 1997). Examples include genes in mothers influencing offspring growth (McAdam and Boutin 2004), and genes in males influencing the date their partner lays a clutch (Brommer and Rattiste 2008). In general, the response to selection in the presence of IGEs is (Bijma and Wade 2008):

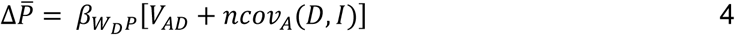

Where *β_W_D_P_* is the selection gradient of an individual’s direct phenotype on fitness, *V_AD_* is the additive direct genetic variance in the trait, *n* is the number of conspecifics an individual interacts with (i.e. group size excluding itself, note this replaces *n−1* used by Bijma and Wade 2008, as they set n as group size *including* the focal individual), and *cov_A_*(*D*,*I*) is the additive genetic covariance between the direct and indirect effects on the trait. The product of *β_W_D_P_* and *V_AD_* is equivalent to the first term in the Price Identity in the absence of an environmental covariance between the trait and fitness (Rausher 1992). The product of *β_W_D_P_* and *ncov_A_*(*D,I*) represents the correlated evolution of the social environment that occurs because of the genetic covariance between an individual’s effect on its own phenotype (direct genetic effect; DGEs) and its effect on the phenotype of others (IGEs). This is the correlated evolution of the social environment, or in other words a non-zero transmission bias. Equation 4 makes clear that, in the presence of covariance between DGEs and IGEs, transmission bias in the Price identity is non-random with respect to selection and clearly cannot be ignored. While transmission bias is often ignored because of an assumption that the environment remains constant, considering genetic variance in social interactions makes clear that in the presence of *cov_A_*(*D,I*) the environment cannot remain constant; the social environment will necessarily evolve as a correlated response to selection. In the extreme example of contests for dominance, the resource for which individuals compete (success in a dyadic contest) is absolutely limited. However, as Cooke *et al.* (1990) observed, directional selection on any resource dependent trait can be counteracted by changes in the competitive environment, so the same IGE-based model can be applied to any trait dependent on contests for limited resources (Frank and Slatkin 1992; Wilson 2014). For instance, Muir *et al.* (2013) conducted an experiment on Japanese quail (*Coturnix japonica*), where they applied artificial selection for body mass, which possesses additive genetic variance. They observed no response to selection over 20 generations, despite the simple expectation that mean body mass would increase over time in response to artificial selection. In quail, however, body mass is a proxy for competitiveness with pen-mates for access to feed. The heaviest quail were, therefore, the ones that supressed the body mass of their pen-mates the most, by outcompeting them for access to feed. As such, by artificially selecting the heaviest individuals, Muir *et al.* were also selecting for those that reduced the body mass of their pen mates the most. As these traits possessed additive genetic variance, the result was the evolution of direct breeding values for body mass, but also the evolution of breeding values for increased suppression of pen-mates’ body masses. Therefore, there were DGEs for body mass, IGEs for the body mass of pen-mates, and a negative DGE-IGE covariance, overall giving no change in mean body mass. A similarly strong negative covariance between direct and indirect genetic variance in performance was found for diameter at breast height in plantations of Eucalyptus trees (*Eucalyptus globulus*), presumably due to competition with neighbouring trees for light or other resources (Costa e Silva et al. 2013). In both these examples the competitive ability of individuals can evolve, but this leads to the evolution of equally more competitive social environments, and so mean of the trait under selection does not change across generations.

## Indirect genetic effects on fitness

If we consider fitness as a trait influenced by social interactions, then conspecifics can influence each other’s fitness following existing IGE models (Bijma 2011):

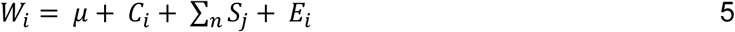

Where individual *i*’s fitness (*W_i_*) depends on the population mean (*μ*), as well as *i*’s direct competitive ability (*C_i_*), the sum of the social effects of its *n* neighbours (*Σ_n_ S_j_*) and an environmental/residual component (*E_i_*; Bijma 2011). This is an analogous framework to the one proposed by Cooke *et al.* (1990), for the evolution of clutch size in birds, subsequently built upon by Frank and Slatkin (1992). This simply says that an individual’s fitness will be influenced by its own competitive ability (e.g. its weapon size) but also by the competitive abilities of other individuals in the group/population (see also models for “social selection”, e.g. Goodnight et al. 1992; Eldakar et al. 2010).

If we wish to consider how these social effects might constrain or facilitate the evolution of fitness, we need to consider the genetic basis of competitive ability and social effects on others’ fitness (following Cooke *et al.* (1990) and Frank and Slatkin 1992). The direct competitive abilities of individuals can be partitioned to an additive genetic component and a non-genetic component. Similarly, an individual’s social effects can be divided into genetic and non-genetic effects on its competitors’ fitness. There is, therefore, additional genetic variance in fitness, stemming from competitors, alongside the more traditionally considered direct genetic variance stemming from the focal individual. This additional genetic variance can contribute to the evolution of fitness. The expected change in mean fitness in the presence of IGEs (when unrelated individuals interact and in the absence of multilevel selection) is given by (note that, as fitness is always maximally selected upon, while the relationship between fitness and fitness passes through zero and is linear, *β_W_D_P_* is at the maximum of 1; Hereford et al. 2004):

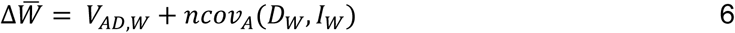

There are two important things to note from eq. 6. First, when *cov_A_*(*D_w_,I_w_*) is 0, we recover the FTNS. This would be true, however, only when there is no intra-specific competition. Instead, often an individual’s fitness gains will necessarily detract at least somewhat from the fitness of others and *cov_A_*(*D_w_,I_w_*) will be negative. A negative *cov_A_*(*D_w_,I_w_*) will reduce the rate of evolution of mean fitness, which we have seen is a result of the evolution of a deteriorating environment. If *cov_A_*(*D_w_,I_w_*) is sufficiently negative, Δ*W̅* can equal 0 despite *V_AD,W_* being non-zero. This will occur when fitness is completely zero-sum, such that any fitness accrued by one individual is equal to the fitness lost by a competitor or competitors (e.g. contests over a limited resource). Therefore, *cov_A_*(*D_w_,I_w_*) represents an explicit measure of the degree to which adaptation will be constrained by competition, thereby counteracting the continual evolution of increased mean fitness as predicted by the FTNS (c.f. Cooke et al. 1990; Frank and Slatkin 1992). *cov_A_*(*D_w_,I_w_*) also represents an explicit modelling of environmental deterioration, and of a form of transmission bias, in terms of the contribution of IGEs (changes in the social environment) to the change in mean fitness. Direct breeding values for fitness are still expected to increase across generations, as selection for fitness always occurs. The effect on fitness at the phenotypic level, however, is counterbalanced by the evolution of an increasingly competitive (deteriorating) environment resulting from IGEs on fitness(Cooke et al. 1990; Frank and Slatkin 1992). The degree to which fitness increases are counterbalanced by a deteriorating social environment, and hence the degree to which fitness is zero-sum is measured by *cov_A_*(*D_w_,I_w_*).

## Evolution without adaptation

While fitness IGEs might constrain the evolution of mean fitness (adaptation), the continued evolution of DGEs on fitness means that traits correlated with fitness DGEs can still evolve (unless these traits are also subject to IGEs; see Box 1). This is analogous to the situation observed by Muir *et al.* discussed above. In Muir *et al.* (2013), body mass could not evolve as it was subject to IGEs, but the competitiveness of individual quail was able to evolve. This commonly occurs in livestock selected for increased yields, when pecking or biting behaviours increase across generations, but yields do not (Ellen et al. 2014). This occurs because traits related to social competition (e.g. aggressive pecking) are correlated with the *direct* additive genetic variance in the yield trait (e.g. body mass). Traits related to social competition can, therefore, increase, while overall performance (e.g. yield) remains constant because of the evolution of more competitive environments. In the case of fitness, traits related to fitness, such as weapon size or the brightness of a sexual display trait, can evolve over time even when mean fitness does not evolve (but see Box 1). This, therefore, solves the apparent problem posed by the two statements we made at the start of this paper. Evolution occurring in populations where mean fitness is not evolving is in fact compatible with Fisher’s FTNS and the PI once IGEs on fitness are considered. Furthermore, evolution without adaptation is absolutely required for the evolution of environmental deterioration to occur (in the form of the evolution of more competitive rivals), yet this is often not made explicit. If traits related to competitive ability cannot evolve then the environment cannot deteriorate in this manner.

Neither the general ideas, nor models that we have outlined here are new. Applying these ideas and models to fitness itself, however, clarifies when evolution and adaptation are expected to occur, and when they are not. Arguably, Fisher would have classified all changes in indirect effects as environmental deterioration, meaning that we should not model them explicitly here. However, as this change has an additive genetic basis and is correlated with changes in fitness due to direct genetic effects, it seems essential to include them in our models for the evolution of fitness. Furthermore, there are additional insights into trait evolution and adaptation that come from considering IGEs on fitness and fitness-related traits.

## The evolution of maladaptation

An interesting outcome of models for evolution in the presence of IGEs is that traits can respond in the *opposite* direction to selection if a negative *cov_A_*(*D,I*) outweighs the influence of direct effects (Griffing 1967; Moore et al. 1997; more formally, when —11(*cov_A_*(*D,I*)) > *V_AD_*/*n*). In these cases, selection favours individuals whose indirect effects reduce the population mean more than their direct effects increase it. What this means for the evolution of fitness is that, although *V_AD,W_* can never be less than zero, Δ*W̅* can be negative (i.e. the evolution of maladaptation), if *cov_A_*(*D_W_,I_W_*) is strong enough (—1(*cov_A_*(*D_W_,I_W_*)) > *V_AD,w_*/*n*; note this is analogous to the possible decrease in mean fitness when selection acts on linked loci (Moran 1963), just that the fitness effects of the loci are observed in different individuals). This is distinct from cases where fitness decreases due to a deterioration in the non-social or abiotic environment, as the change in fitness caused by evolution of IGEs is the direct result of selection (effectively for individuals that supress others the most). Such an effect has been observed in populations of flour beetles (*Tribolium castaneum*), where artificial selection for individuals with *increased* reproductive output caused the mean reproductive output across the populations to *decrease* over time (Wade 1976). This may apply more generally to populations that are approaching or above a habitat’s carrying capacity, and so mean fitness is expected to decline in subsequent generations. That the FTNS only ever allowed for an increase in fitness (adaptation, but not maladaptation) has been one of its major criticisms (Frank and Slatkin 1992). Modelling the evolution of fitness in the presence of IGEs allows maladaptation to occur, reconciling the FTNS with empirical observations.

## Indirect genetic effects and density dependence

Including IGEs in the expected change in mean fitness also leads to useful links between quantitative genetics and population biology. For instance, eq. 6 takes similar form to the logistic model of density-dependent per capita population growth:

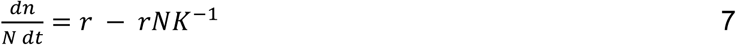

In the logistic model the rate of per capita population growth 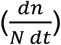 is positively affected by the intrinsic rate of increase of the population (r), while -*rNK^−1^* represents the degree to which per capita population growth is reduced by per capita increases in death rates and decreases in birth rates as the population approaches its carrying capacity (*K*). Such density dependence results from social interactions (such as competition for space or food) among individuals that cause them to supress the birth rate or increase the death rate of others. These social effects may well have a genetic component, and hence be IGEs. When populations are far below *K*, indirect effects on fitness are expected to be relatively weak. In this scenario *V_AD,W_* can exceed *ncov_A_*(*D_w_,I_w_*) and mean fitness can evolve. This is analogous to r-selection, as a low contribution from *ncov_A_*(*D_w_, I_w_*) due to non-limiting resources allows the evolution of fitness and so rapid population growth. However, as the population size approaches *K*, negative social effects on fitness become stronger, and *ncov_A_*(*D_w_,I_w_*) will eventually be large enough to equal *V_AD,W_*, and mean fitness can no longer evolve. The change in mean fitness may even reduce below zero, causing the population size to return below *K*.

Density-dependent selection has typically been modelled from a framework where genotypes differ in their sensitivity to competition, which has led to the prediction of the evolution of increased carrying capacity at high density (an increase in “efficiency” of organisms; MacArthur 1962). The model including IGEs on fitness, however, makes an additional prediction: at high density, we expect the evolution of increased ability to depress the survival and reproduction of others as the population approaches carrying capacity (in Fisher’s words: “life is made somewhat harder to each individual when the population is larger”; Fisher 1930). This process ought to result in the evolution of *reduced* K. It is not currently clear the degree to which density dependent selection in nature favours increased efficiency versus enhanced ability to supress the fitness of others.

It is tempting to directly relate the group size, *n*, in eq. 6 with the population size, *N*, in eq. 7, but these are not necessarily equivalent. All individuals within a population are unlikely to interact with one another socially to the degree that they might depress one another’s fitness, so if population size (*N*) increases but density does not (i.e. the population expands into uninhabited space) then the number of socially interacting individuals (*n*) will not change. It is also generally expected that larger groups sizes should weaken *cov_A_*(*D_W_,I_W_*), as more distant or more weakly interacting individuals who do not influence each other’s fitness are included within progressively larger groups (Fig. 1, top panel, see also Bijma 2010b). If, however increasing population size implies greater *density*, as well as simply more individuals, then social interactions may well get more intense (Fig. 1, bottom panel). This would imply a greater, or at least stationary, *cov_A_*(*D_W_,I_W_*) as *n* increases, and so the product *cov_A_*(*D_W_,I_W_*) would contribute increasingly to Δ*W̅*. The explicit inclusion of IGEs on fitness, therefore, results in the emergence of density-dependent per capita reproduction through social effects.

**Figure 1.**
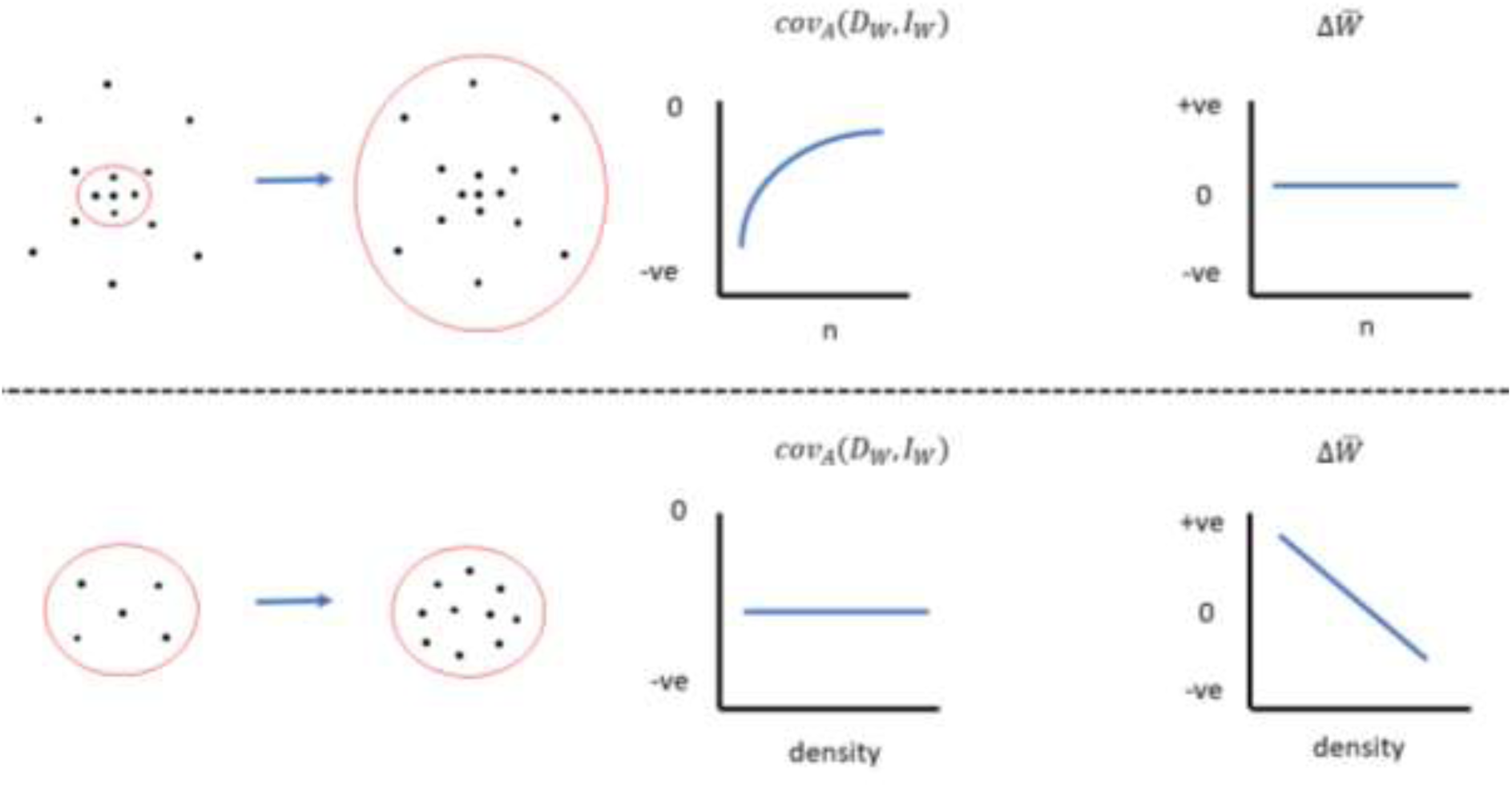
The relationship between *n*, density, *cov_A_*(*D_w_,I_w_*), and expectations for Δ*W̅*. Here we assume that the fitness of individuals is based on competition for limited resources, and so *cov_A_*(*D_w_,I_w_*) ranges from 0 to strongly negative. If we simply increase the number of individuals considered (top panel), then we expect *cov_A_*(*D_w_,I_w_*) to approach 0, as the additional individuals are less closely associating with each other, decreasing the mean social effect individuals have on each other. This balances the increase in *n*, giving a stationary Δ*W̅*. Here we have depicted Δ*W̅* remaining at 0, assuming the population has reached a point that resources are completely preventing further evolution of increased reproduction. If, however, we increase the density of the individuals, as well as their number (bottom panel), then the *cov_A_*(*D_w_,I_w_*) may be stationary, or even become more negative, as the number of individuals increases. This reduces Δ*W̅*, in our example from an initial period of increasing fitness (below *K*), through no change (at *K*) and then to a decline (above *K*). This is the emergence of density dependent reproduction, only apparent through the FTNS when IGEs for fitness are considered.

The magnitude of the reduction in Δ*W̅* caused by a negative *cov_A_*(*D_W_,I_W_*) depends on how completely mean fitness in the population is constrained. Mild constraints will mean a *cov_A_*(*D_W_,I_W_*) closer to zero (but still negative), and therefore a reduced, but not completely eliminated, increase in mean fitness across generations. Absolute constraints mean a strong negative *cov_A_*(*D_W_,I_W_*), and no change in mean fitness (no adaptation) or even a decrease (maladaptation). Therefore, the difference between *V_AD,W_* and *cov_A_*(*D_W_,I_W_*) is a measure of the magnitude of the constraints on the evolution of mean fitness. How *cov_A_*(*D_W_,I_W_*) changes with *n* is an indication of the strength of density dependence, but cannot be predicted beforehand. This instead remains an empirical question to be answered. *cov_A_*(*D_W_,I_W_*) can be converted to a correlation between an individual’s direct and indirect genetic effects on fitness to compare across populations, with 0 indicating no constraints and −1 indicating complete constraints, as found when analysing the evolution of dominance contests (Wilson et al. 2009, 2011; Sartori and Mantovani 2013). Positive values would indicate synergistic effects such as Allee effects (Allee 1931). In terms of hard and soft selection, a correlation of 0 would indicate that selection is hard (not dependent on the traits of others and leads to adaptation) while a correlation of −1 would indicate that selection is completely soft (entirely dependent on the trait of an individual relative to others and does not lead to adaptation).

## Adaptation when direct genetic variance in fitness is zero

A final outcome of considering IGEs on fitness is that fitness can evolve (adaptation or maladaptation can occur) in populations where direct genetic variance in fitness is zero (*V_AD,W_* = 0), if there are IGEs on the fitness of *related* conspecifics. When unrelated individuals interact, if *V_AD,W_* is zero, *cov_A_*(*D_w_,I_w_*) is then undefined and, following eq. 6, Δ*W̅* is zero. However, if related individuals interact, the expected change in mean fitness follows (and Wade 2008):

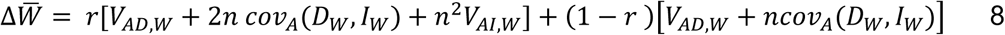

Where *r* is the mean coefficient of relatedness between interacting individuals, *V_AI,W_* is the additive indirect genetic variance for fitness, and other terms are as defined for eq. 6. This allows a change in mean fitness when *V_A_D_,W_* and *cov_A_*(*D_W_,I_W_*) = 0, as long as *V_AI,W_>* 0 *and r ≠* 0:

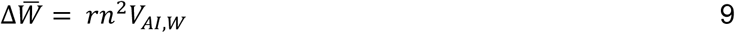

So, in contrast with a simple interpretation of FTNS, population mean fitness can evolve even in the absence direct genetic variance in fitness, as long as fitness-relevant social interactions are with relatives and there are IGEs for fitness. Note in these equations for the response to selection in the presence of IGEs, *r* can be replaced without altering the equations by g, the relative strength of multilevel selection (Bijma and Wade 2008). As such, the presence of multilevel selection can also allow adaptation (or maladaptation) to occur when *V_A_D_,W_* is zero, as long as *V_AI,W_*> 0 and *g* ≠ 0 (see Bijma and Wade 2008 for when both *r* and *g* are non-zero, and see also McGlothlin et al. 2010).

Given that populations in equilibrium conditions are typically expected to show very little *V_AD,W_* (Fisher 1930), this provides a mechanism for those populations to still adapt. For instance, in a population of North American red squirrels (*Tamiasciurus hudsonicus*) *V_AD,W_* was found to be essentially zero, but maternal genetic effects on fitness were present (McFarlane et al. 2015). Maternal genetic effects are a specific form of IGE where a mother’s genes (e.g. for milk production) influence the traits of her offspring. When parents interact with offspring, *r* is non-zero. Models for evolution in the presence of maternal genetic effects are then valid, which allows the population to evolve, albeit with a lag due to the crossgenerational effect (Kirkpatrick and Lande 1989; Mousseau and Fox 1998). Therefore, fitness can change from generation to generation, despite lacking direct additive genetic variance. This is not a new result, as evolution and adaptation in the presence of maternal genetic effects and IGEs in general is accepted. Worth noting is that, as direct breeding values for fitness are not changing across populations, the breeding values for any traits genetically correlated with these will also not change. A trait may evolve, however, if it is genetically correlated with indirect breeding values for fitness.

## Conclusions

Fig. 2 illustrates four situations which correspond to our formulation for the change in mean fitness we have outlined above (although we do not plot the case where DGEs for fitness are absent but IGEs among relatives and/or in the presence of multilevel selection do occur, see the section on “Adaptation when direct genetic variance in fitness is zero”). These represent a complete range of cases: when DGEs for fitness are either absent or present, when IGEs are either absent or present, and if both DGEs and IGEs are present, if they positively or negatively covary. We indicate the consequences each situation has for the expected evolution of mean fitness (adaptation), as well as for the evolution of other traits within the population (evolution by natural selection). These demonstrate that considering the evolution of fitness as the response to selection in the presence of IGEs allows us to account for many situations observed in nature and captive breeding. Frank and Slatkin stated that “fitness…increases by an exact amount because of natural selection but simultaneously increases or decreases by an unpredictable amount because of the environment”(Frank and Slatkin 1992). We hope we have shown here that, by incorporating IGEs into our models, a portion of this change caused by the environment is predictable.

**Figure 2a-d.**
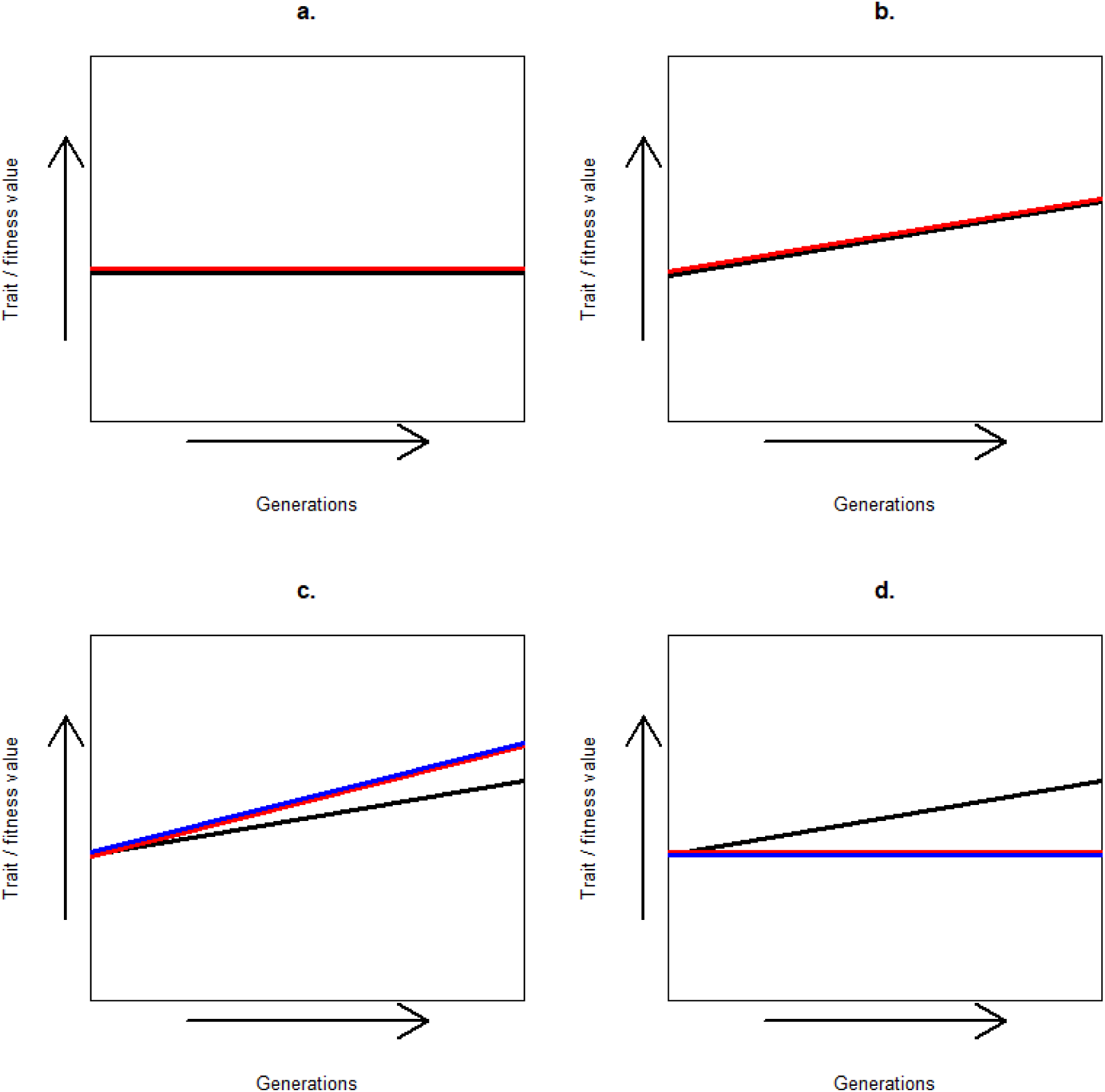
How fitness (red) and a trait (black) are expected to change across generations. Note the scale for both the trait and fitness is arbitrary; we do not necessarily expect a trait and fitness to increase at exactly the same rate in scenario b. for example. For simplicity we assume that interactions are with non-relatives (*r* = 0) and there is no multilevel selection (*g* = 0). a: No DGEs for fitness, no IGEs. No genetic variance in fitness. Neither adaptation nor any evolution will occur. b: DGEs for fitness, but no IGEs. Heritable variance in fitness is present, and so mean fitness is expected to evolve over time in line with the FTNS. Traits genetically correlated with fitness are also able to evolve. Both adaptation and evolution can occur. c: DGEs and IGEs for fitness, positive DGE-IGE covariance. Heritable variance in fitness is present, and so mean fitness is expected to increase over time, and rapidly as the positive DGE-IGE covariance shifts the response in the same direction as selection. Traits genetically correlated with fitness will evolve, although only as fast as fitness if they too are influenced by IGEs (blue line). Evolution and rapid adaptation. d: DGEs and IGEs for fitness, negative DGE-IGE covariance for fitness. The expected evolution of fitness will be reduced, possibly to zero or even below. However, as direct breeding values for fitness will still be increasing across generations, traits genetically correlated with fitness may evolve, unless they too are influence by IGEs (blue line). This corresponds to situations where livestock under artificial selection for increased yield have shown no evolution of yield but do show increases in aggressive behaviours such as biting or pecking, as well as the instances of sexual selection described in the text. Evolution but no adaptation.

In summary, considering IGEs on fitness allows us to reconcile the FTNS and the PI with several observations: 1) it allows evolution even when adaptation is not occurring. This was acknowledged by Fisher, and is implied by models for trait evolution in the presence of IGEs, but appears impossible under conventional understandings of the FTNS and PI. 2) It allows the evolution of maladaptation, reconciling the FTNS with empirical observations. 3) Including *n* in the equation for the change in mean fitness reveals density-dependence, helping to link quantitative genetics to population biology. 4) It indicates when adaptation can occur even when direct genetic variance in fitness is lacking. Considering IGEs on fitness explicitly models the deterioration of the social environment, a type of transmission bias, and so clarifies how both the evolution of traits and the adaptation of populations is expected to proceed.

## Acknowledgements

We thank Cortland Griswold, Loeske Kruuk, Alastair Wilson and Piter Bijma for comments and discussions that greatly improved this manuscript. We have no conflicts of interest.

## Author contributions

Both authors conceived of the research question, drafted the manuscript, and approved the final version. DNF made the figures.

